# Global repression of mesenchymal genes in epithelial cells by the histone demethylase LSD1

**DOI:** 10.64898/2026.11.04.764245

**Authors:** Robert E. Turnbull, Oksana Gonchar, Kristupas Sirvydis, Mariam Iftikhar, Megan E. Coulson, So Yeon Kwon, James T. Hodgkinson, Paul Badenhorst, Shaun M. Cowley, John W.R. Schwabe

## Abstract

Epithelial-mesenchymal transition (EMT) is an essential process in development, tissue repair, and tumour metastasis, and is triggered by SNAI, ZEB and TWIST transcription factors, that recruit lysine-specific demethylase 1 (LSD1) to repress epithelial genes such as E-cadherin. In mesenchymal cells, LSD1 inhibition restores E-cadherin expression and reduces migration and invasion, consistent with LSD1 inhibition reverting EMT, and offering a therapeutic modality for EMT-associated disease. Recent data, however, suggest context-dependent roles for LSD1, as its inhibition in epithelial cells increases mesenchymal marker expression, potentially promoting EMT. To investigate this discrepancy, we mapped genome-wide binding of LSD1 in HCT116 epithelial cells and used rapid degron-mediated ablation of endogenous LSD1 to measure changes in nascent and steady-state transcription and chromatin accessibility. In epithelial cells, LSD1 binds constitutively-accessible, primed mesenchymal promoters. These are rapidly derepressed after LSD1 ablation, with activation of SNAI2 and ZEB1, which repress epithelial hallmarks such as E-cadherin. Re-introduction of LSD1 downregulates SNAI2 and ZEB1 but, surprisingly, further represses E-cadherin, indicating sustained loss of epithelial characteristics. Collectively, our results show that LSD1 maintains epithelial identity by repressing mesenchymal genes, questioning the therapeutic utility of LSD1 inhibition for EMT control.

## Introduction

Epithelial-to-mesenchymal transition (EMT) and the reverse mesenchymal-to-epithelial transition (MET) play critical roles in morphogenesis and in pathologies including cancer and chronic inflammation. EMT is classified into three types: type 1, mesoendoderm development; type 2, wound healing and fibrosis; and type 3, cancer metastasis (Huang et al. 2022; Zhang et al. 2025). Normal epithelial cells are immotile and maintain tissue integrity through tight junctions and close interactions with the extracellular matrix (ECM). Mesenchyme-inducing factors trigger epithelial junction breakdown, ECM degradation and actin cytoskeleton reorganisation, producing a motile, fibroblast-like morphology (Zhang et al. 2025).

EMT is governed by multiple intersecting pathways. The best-studied initiating signals are the Transforming Growth Factor-β (TGF-β) and Fibroblast Growth Factor (FGF) pathways (Strutz et al. 2002; Kasai et al. 2005), but Wnt (Qi et al. 2014), Hedgehog (Omenetti et al. 2008), and Notch (Timmerman et al. 2004) signalling also contribute significantly. These inputs often converge to activate SMAD proteins, particularly SMAD3 and SMAD4, whose depletion reduces EMT (Hua et al. 2020). This interplay places EMT on a continuum of cell states rather than a binary switch, allowing cells to transition between epithelial and mesenchymal states or to occupy intermediate states (Bakir et al. 2020).

Following EMT induction, upregulated mesenchymal transcription factors (TFs), including SNAI1, SNAI2, TWIST and ZEB1/2, promote expression of mesenchymal markers such as Vimentin (VIM) and N-cadherin and repress epithelial genes such as E-cadherin by binding E-box sequences in their promoters (Wels et al. 2011; Saitoh 2023). SNAI1/2 (Lin et al. 2010a; Lin et al. 2010b) and ZEB1/2 (Shi et al. 2003) interact with and recruit the CoREST complex, leading to repression of epithelial genes (Lin et al. 2010a; Lin et al. 2010b).

CoREST comprises the catalytic subunits lysine-specific demethylase 1 (LSD1) and histone deacetylase 1/2 (HDAC1/2), and a co-repressor protein, RCOR1/2/3 (Hakimi et al. 2002; Shi et al. 2004). CoREST recruitment during EMT-associated transcriptional reprogramming is therefore accompanied by epigenetic modifications (McDonald et al. 2011). HDAC1/2 deacetylates Histone 3 lysine 9 (H3K9ac) to repress gene expression (Lee et al. 2005; Wu et al. 2018), whereas LSD1 can demethylate H3K4me1/2 to repress, or H3K9me1/2 to activate, gene expression (Shi et al. 2004; Lee et al. 2005; Metzger et al. 2005).

The importance of CoREST in driving and maintaining the mesenchymal phenotype is well documented. Elevated LSD1 is associated with metastasis and aggressiveness in several cancers (Metzger et al. 2005; Schulte et al. 2009; Lv et al. 2012; Li et al. 2016; Xie et al. 2020; Bottner et al. 2022). Conversely, LSD1 inhibition or knockdown in metastatic cancer cell lines increases epithelial and represses mesenchymal gene expression, reducing migration and invasiveness (Ferrari-Amorotti et al. 2013; Ferrari-Amorotti et al. 2014; Wang et al. 2015; Xie et al. 2020; Wang and Liu 2023). In somatic epidermal cells, LSD1 inhibition promotes differentiation and suppresses squamous cell carcinoma progression (Egolf et al. 2019). LSD1 has consequently been prioritised as a cancer therapeutic target, with at least nine inhibitors in phase 1 or 2 clinical trials for haematological and solid tumours (Noce et al. 2023).

The role of CoREST in epithelial cells is less clear, but some evidence suggests it may have contradictory roles in maintaining the epithelial cell state. LSD1 depletion in epithelial-like breast cancer cells reduces E-cadherin expression and increases invasion and migration (Hu et al. 2019), and chemical inhibition of LSD1 in OM1 oral squamous carcinoma cells increases ZEB1 and VIM expression (Yamakado et al. 2023). Although the mechanisms remain unclear, the demethylase UTX and the metastasis repressor BRMS1 have been suggested to recruit LSD1/CoREST to a subset of mesenchymal promoters in epithelial cells to mediate repression (Choi et al. 2015; Qiu et al. 2018).

Consequently, although LSD1 inhibition is well established to promote MET in mesenchymal cells, it may prime or induce EMT in epithelial cells challenging therapeutic strategies that rely on LSD1 inhibitors to regulate EMT. To understand the mechanistic basis of these opposing effects, we integrated genome-wide LSD1 occupancy with both nascent and steady-state transcription and chromatin accessibility profiling in epithelial cells following acute LSD1 depletion. This approach has allowed us to resolve cell-state-specific roles for LSD1 in maintaining epithelial identity and regulating mesenchymal cell programmes.

## Results

### Time-resolved, rapid knockdown of LSD1 reveals both immediate and delayed changes in gene expression in epithelial cells

To study the LSD1 regulome in epithelial cells we used HCT116 cells, an epithelial colorectal carcinoma line widely used to model EMT. Both copies of the endogenous LSD1 locus were CRISPR-engineered to carry the FKBP12^F36V^ degron (for depletion) and a FLAG tag (for visualisation) (Turnbull et al. 2025); NAR in press). Treatment of LSD1-FKBP12^F36V^-FLAG cells with the PROTAC dTAG^V^-1 depleted >95% of LSD1 within 1 h (Supplementary Fig. S1A). LSD1 depletion also reduces the CoREST subunit RCOR1 (Turnbull et al. 2025; NAR in press), suggesting the complex is no longer functional. Coupling depletion with nascent and steady-state transcriptomics identified direct LSD1 targets, and timed dTAG^V^-1 washout allowed us to test whether transcriptional changes were stable or reversible on LSD1 re-introduction.

We first performed mRNA-sequencing on control and dTAG^V^-1-treated cells at 6, 12, 24 and 48 h to monitor rapid and sustained changes in expression. Principal component analysis (PCA) clearly separated control and treated triplicates at all timepoints (Supplementary Fig. S1B). After filtering for genes with read counts above 50 in at least three samples, differential expression analysis (pAdj < 0.05, fold change ≥ 1.4) revealed similar numbers of differentially expressed genes (DEGs) at all four timepoints (Fig. 1A). Strikingly, over 95% of DEGs at 6 h were upregulated, with only 23 genes downregulated. Downregulated DEGs increased to 52 at 48 h (Fig. 1A) but remained a minor proportion at all timepoints and showed little overlap between timepoints (Supplementary Fig. S1C). This is consistent with LSD1 acting primarily as a repressor through H3K4me1/2 demethylation and/or CoREST-associated HDAC deacetylation of H3K9.

**Figure 1.**
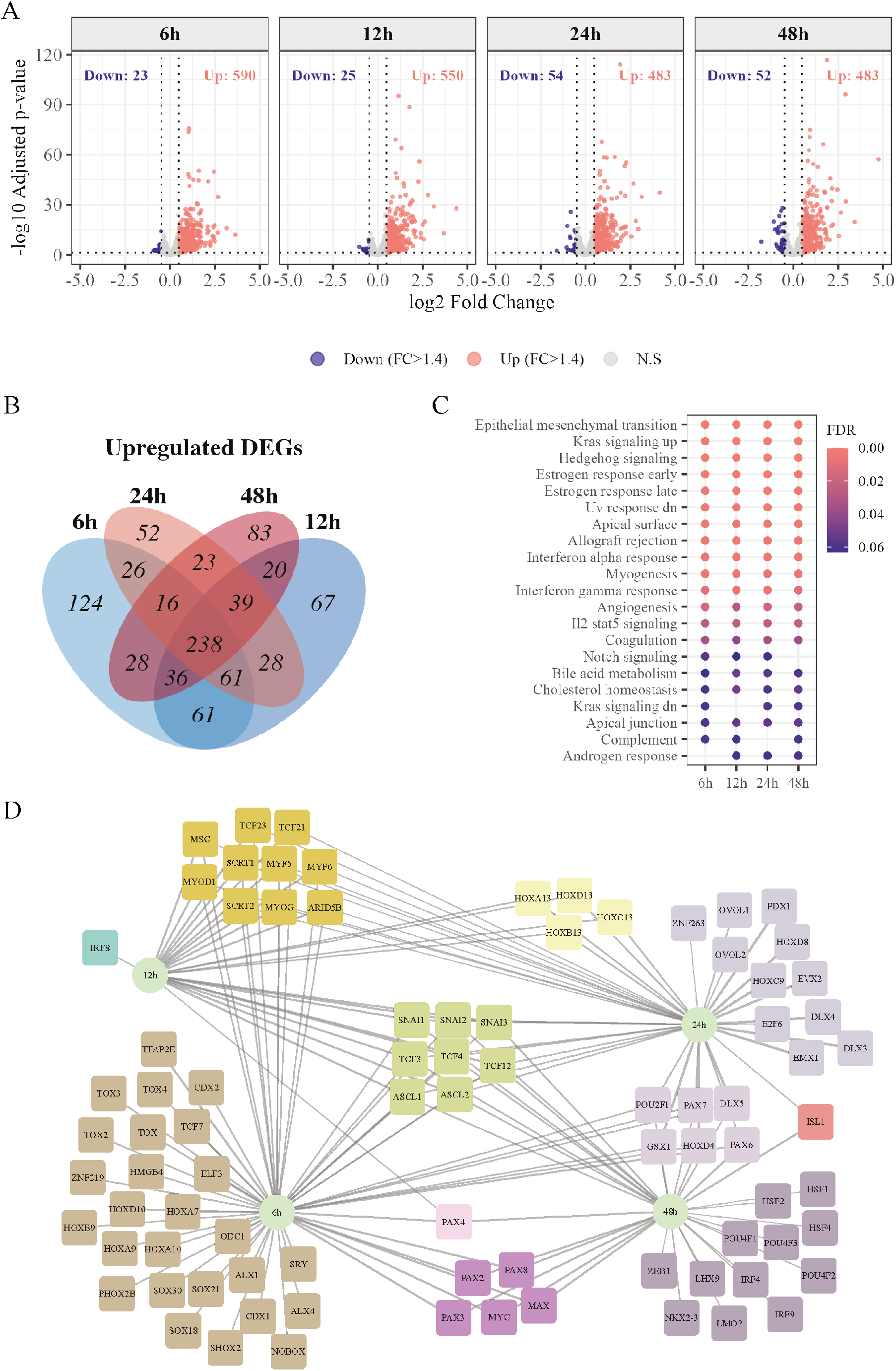
LSD1 depletion causes rapid, sustained changes in gene expression. (A) Volcano plots of significantly down-(blue) and upregulated (red) genes following LSD1 depletion over 48 h. Horizontal dashed line, pAdj ≤ 0.05; vertical dashed lines, |log2FC| ≥ 0.485 (1.4-fold change). (B) Venn diagram of upregulated DEGs at 6, 12, 24 and 48 h. (C) Dot plot of significantly perturbed pathways identified by GSEA at each timepoint. (D) Network map of TFs with motifs identified by iRegulon (Janky et al. 2014) in upregulated DEGs at each timepoint.

In contrast, upregulated DEGs overlapped substantially over the time course, with 390 mRNAs shared between three or more timepoints, and 54% of genes upregulated at 6 h were also upregulated at 48 h, indicating rapid and sustained derepression of a transcriptional programme (Fig. 1B). Multiple independently derived HCT116 LSD1-FKBP12^F36V^-FLAG clones (Supplementary Fig. S1A, D, E) showed essentially identical clustering by PCA 24 h after LSD1 depletion (Supplementary Fig. S2A), with 297 overlapping upregulated DEGs (Supplementary Fig. S2B) with well-correlated fold changes (Supplementary Fig. S2C) and similar trends in gene expression (Supplementary Fig. S2D), demonstrating high reproducibility. Downregulated DEGs by contrast, overlapped little between clones (Supplementary Fig. S2B), suggesting these changes are non-specific and further confirming that LSD1 primarily represses gene expression in HCT116 cells.

### LSD1 ablation activates EMT pathway genes with DNA-binding motifs for known TF drivers of EMT

HCT116 cells are epithelial but can undergo EMT in response to several treatments (Liu et al. 2017; Kim et al. 2019b; Xu et al. 2022), providing a useful EMT model and for defining mechanisms ensuring stability of epithelial state. Gene set enrichment analysis (GSEA) of all expressed genes after filtering showed that EMT-associated genes were the most significantly enriched at every timepoint following LSD1 depletion (Fig. 1C). Analysis of TF motifs within 500 bp of the transcription start sites (TSSs) of upregulated DEGs by iRegulon identified consensus sites for several well-characterised EMT-promoting TFs (Fig. 1D): motifs for the hallmark mesenchymal TFs SNAI1, SNAI2, SNAI3, TCF3 and TCF12 were found in upregulated genes from 6 to 48 h, and ZEB1 motifs at 48 h. Although many upregulated genes were shared between timepoints, distinct motifs were also identified, with the most unique motifs at 6 h. This suggests that LSD1 depletion relieves repression of a network of EMT-related genes initially driven by different TFs at 6 h than at 12–48 h.

### Repression of EMT-related genes is mediated through direct binding of LSD1

Overlap of upregulated DEGs with an EMT gene list curated from the EMTome (http://ec2-3-231-76-84.compute-1.amazonaws.com/emtome/index.html) showed that ∼33% at each timepoint were associated with promoting EMT (Supplementary Fig. S3A-D), with 82 upregulated at all timepoints (Fig. 2A). Our previous LSD1 ChIP-seq data (GSE330806; GSM9733045-48) showed that most upregulated mesenchymal DEGs were LSD1-bound and likely direct targets: 87% at 6 h, decreasing slightly to 79% at 48 h, possibly reflecting increasing secondary responses (Fig. 2B). The same trend was observed for all upregulated DEGs (Supplementary Fig. S3E). Of the 82 mesenchymal genes shared between timepoints, 70 were bound by LSD1 at the promoter (Fig. 2C; Supplementary Fig. S3F). These included the hallmark mesenchymal genes *SNAI2*, *ZEB1* and *VIM*, and strong EMT drivers such as *CHN1* (Zhao et al. 2021), *FGF2* (Strutz et al. 2002), *FGF9* (Chang et al. 2021), *PLAU* (Chen et al. 2021), and its receptor *PLAUR* (Qin et al. 2024) (Fig. 2C). These results suggest that upregulation of mesenchymal genes following LSD1 depletion results from loss of LSD1 from promoter.

**Figure 2.**
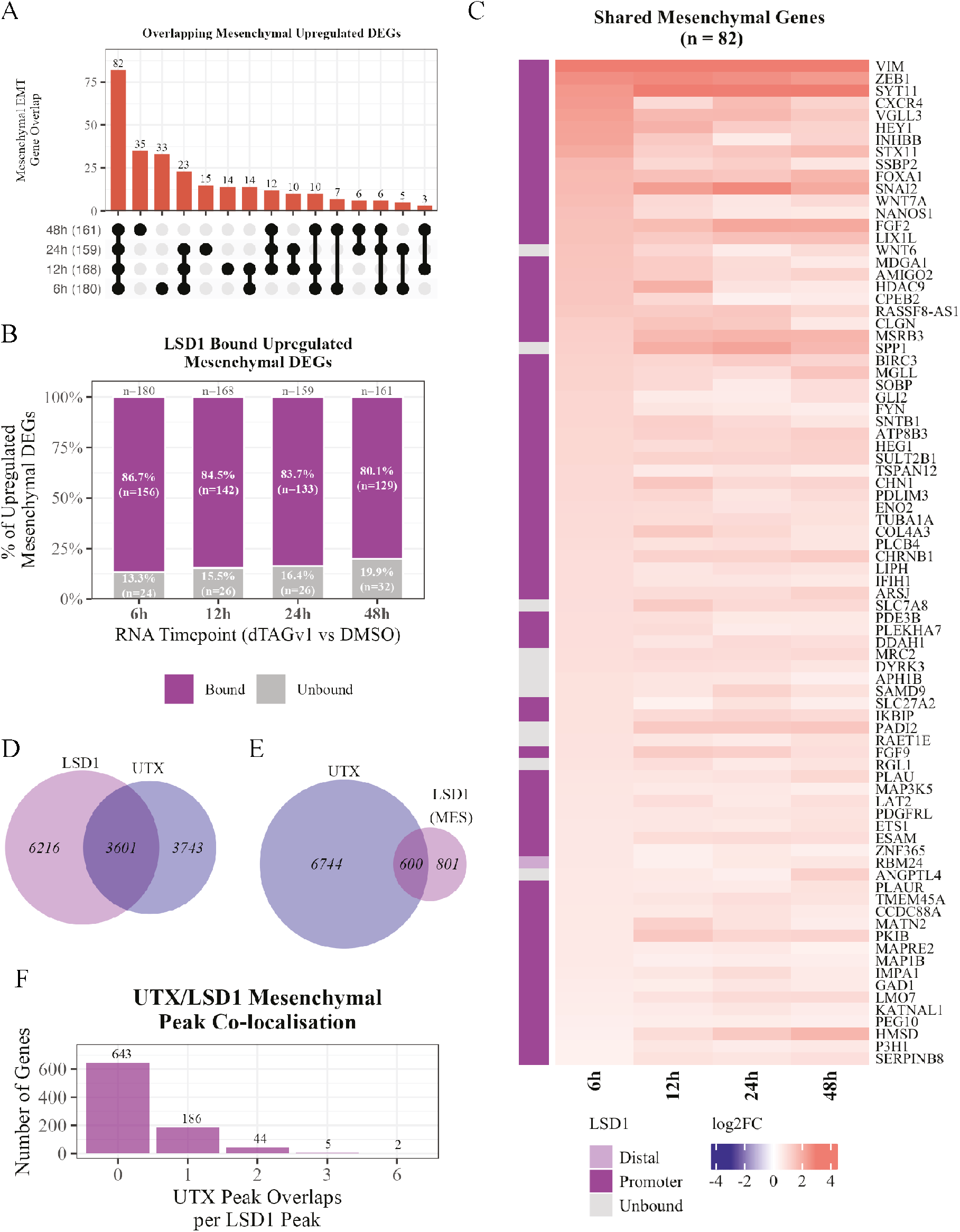
LSD1 depletion results in upregulation of mesenchymal genes. (A) UpSet plot of upregulated mesenchymal DEGs at each timepoint. Connected black circles identify overlapping groups; overlaps were computed in “distinct” mode, i.e. each gene belongs to only one overlap. (B) Stacked bar plot of the percentage of upregulated mesenchymal DEGs at each timepoint directly bound by LSD1. (C) Heatmap of log2 fold change of the 82 mesenchymal DEGs upregulated at every timepoint, with LSD1 binding at the promoter, at a distal site or unbound. (D, E) Venn diagrams of LSD1-bound genes (D) or LSD1-bound mesenchymal genes (E) in HCT116 cells and UTX-bound genes in HeLa cells. (F) Bar plot of the number of UTX peaks directly overlapping LSD1 peaks at mesenchymal genes in HCT116 cells.

LSD1-mediated repression of several mesenchymal genes has been associated with recruitment by BRMS1 and UTX (Choi et al. 2015; Qiu et al. 2018). Comparing our LSD1 ChIP-seq data with a public UTX ChIP-seq dataset from HeLa cells (Kagda et al. 2025) (https://www.encodeproject.org/; ENCFF371TDC) showed that 37% of all LSD1-bound genes and 43% of LSD1-bound mesenchymal genes also had UTX binding sites (Fig. 2D, E). Within the 600 overlapping mesenchymal genes, 237 of 880 (27%) LSD1 binding sites directly overlapped UTX binding (Fig. 2F). Although the datasets derive from different cell lines, this suggests that global repression of mesenchymal genes by LSD1 may be mediated, at least in part, through recruitment by UTX.

### LSD1 ablation increases RNA Pol II loading and transcription

To determine whether increased steady-state mRNA levels reflect increased transcription through release of promoter-proximally paused RNA polymerase II (Pol II), we performed nascent RNA-sequencing (qPRO-seq). As with mRNA-seq, many more genes showed increased than decreased nascent transcription at both 2 h and 6 h after dTAG^V^-1 treatment (Fig. 3A), with 447 genes upregulated at both timepoints and little overlap between downregulated genes (Supplementary Fig. S4A, B). Interestingly, 356 genes showed higher nascent transcription at 2 h than at 6 h, suggesting an initial, transient burst of transcription that diminishes by 6 h.

**Figure 3.**
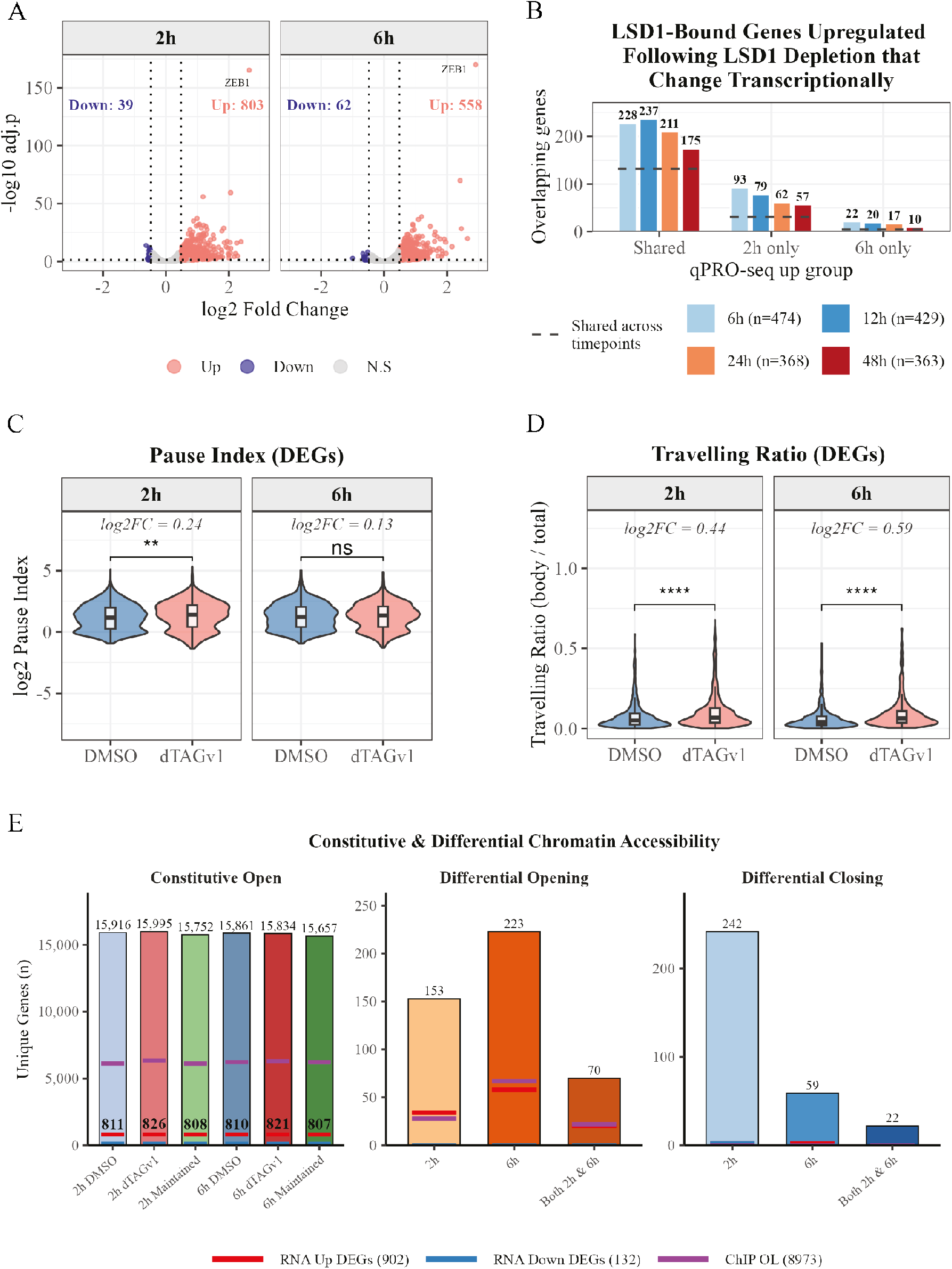
LSD1 depletion increases RNA polymerase loading at gene promoters within open chromatin. (A) Volcano plots of genes with significantly decreased (blue) or increased (red) transcription at 2 h and 6 h after LSD1 depletion. Horizontal dashed line, pAdj ≤ 0.05; vertical dashed lines, |log2FC| ≥ 0.485 (1.4-fold change). (B) Bar plot of LSD1-bound upregulated mRNA DEGs at each timepoint that show transcriptional changes at both 2 h and 6 h (shared), only 2 h or only 6 h. Horizontal dotted lines indicate the number of genes shared across all timepoints. (C) Violin plot of the pause index (reads within ±300 bp of the TSS divided by gene-body reads) and (D) travelling ratio (gene-body reads divided by total reads) of all genes with significantly increased transcription at 2 h and 6 h. (E) Bar plots of constitutive and differential chromatin accessibility: constitutively open regions in control and LSD1-depleted cells and those maintained in both (left), and regions showing differential opening (middle) or closing (right) at 2 h and 6 h after LSD1 depletion. ns, not significant; **p ≤ 0.01; ****p ≤ 0.0001.

Genes with increased nascent transcription at both 2 h and 6 h showed clear overlap with LSD1-bound genes upregulated at the mRNA level at 6 h (48%), 12 h (55%), 24 h (57%) and 48 h (48%) (Fig. 3B). Genes with increased nascent transcription only at 2 h accounted for 20% of upregulated mRNAs at 6 h, decreasing to 16% at 48 h, whereas genes increased only at 6 h accounted for merely 5% (Fig. 3B). LSD1-bound mesenchymal genes upregulated at the mRNA level showed a similar trend (Supplementary Fig. S4C). This suggests nascent transcriptional changes at both 2 h and 6 h reflect increased mRNA expression at 6, 12 and 24 h after LSD1 depletion, while changes seen only at 2 h correlate with increased mRNA at earlier timepoints.

Transcriptionally upregulated genes showed a significant ∼1.2-fold increase in pause index at 2 h after LSD1 depletion, which normalised by 6 h (Fig. 3C), and a travelling ratio that increased 1.35-fold at 2 h and 1.5-fold at 6 h (Fig. 3D). These increases were not observed across all expressed genes (Supplementary Fig. S4D-E) but only for those upregulated after LSD1 depletion. This suggests LSD1 depletion increases Pol II loading at promoters: at 2 h promoter-proximal accumulation elevates pausing, and by 6 h increased gene-body transcription normalises the pause index.

### LSD1 depletion does not result in extensive changes in chromatin accessibility

To determine whether increased promoter loading, transcription and mRNA expression correlated with chromatin opening, we performed ATAC-seq at 2 h and 6 h. We identified >15,000 constitutively open promoter regions (within 500 bp of a TSS) that were largely maintained between control and dTAG^V^-1 treatment (Fig. 3E). Strikingly, 6,000 (>60%) LSD1 binding sites and 90% of upregulated genes across all timepoints were associated with constitutively open chromatin (Fig. 3E). LSD1 depletion significantly opened 153 regions at 2 h, increasing to 223 at 6 h, whereas closing regions were more abundant at 2 h (242) than at 6 h (59) (Fig. 3E). Neither opening nor closing regions overlapped substantially with LSD1 binding or mRNA changes (Fig. 3E).

Together, these data suggest that LSD1 occupies constitutively open chromatin regions within gene promoters that are primed for transcription initiation but repressed by LSD1 activity. When LSD1 is depleted, this repression is lost, allowing increased Pol II loading and transcription.

### LSD1 depletion results in a partial and reversible EMT phenotype

Mesenchymal gene upregulation suggests that LSD1 maintains epithelial identity in HCT116 cells, analogous to its accepted role in mesenchymal cells. To test effects on protein expression, we examined the hallmark proteins SNAI1, SNAI2, VIM, ZEB1 and E-cadherin. SNAI2, ZEB1 and VIM increased significantly 24 h after LSD1 depletion, whereas the epithelial structural protein E-cadherin decreased over the time course (Fig. 4A; Supplementary Fig. S5A). SNAI1 decreased over the time course, as its stability depends on interaction with LSD1 (Zhou et al. 2004; Lin et al. 2010b).

**Figure 4.**
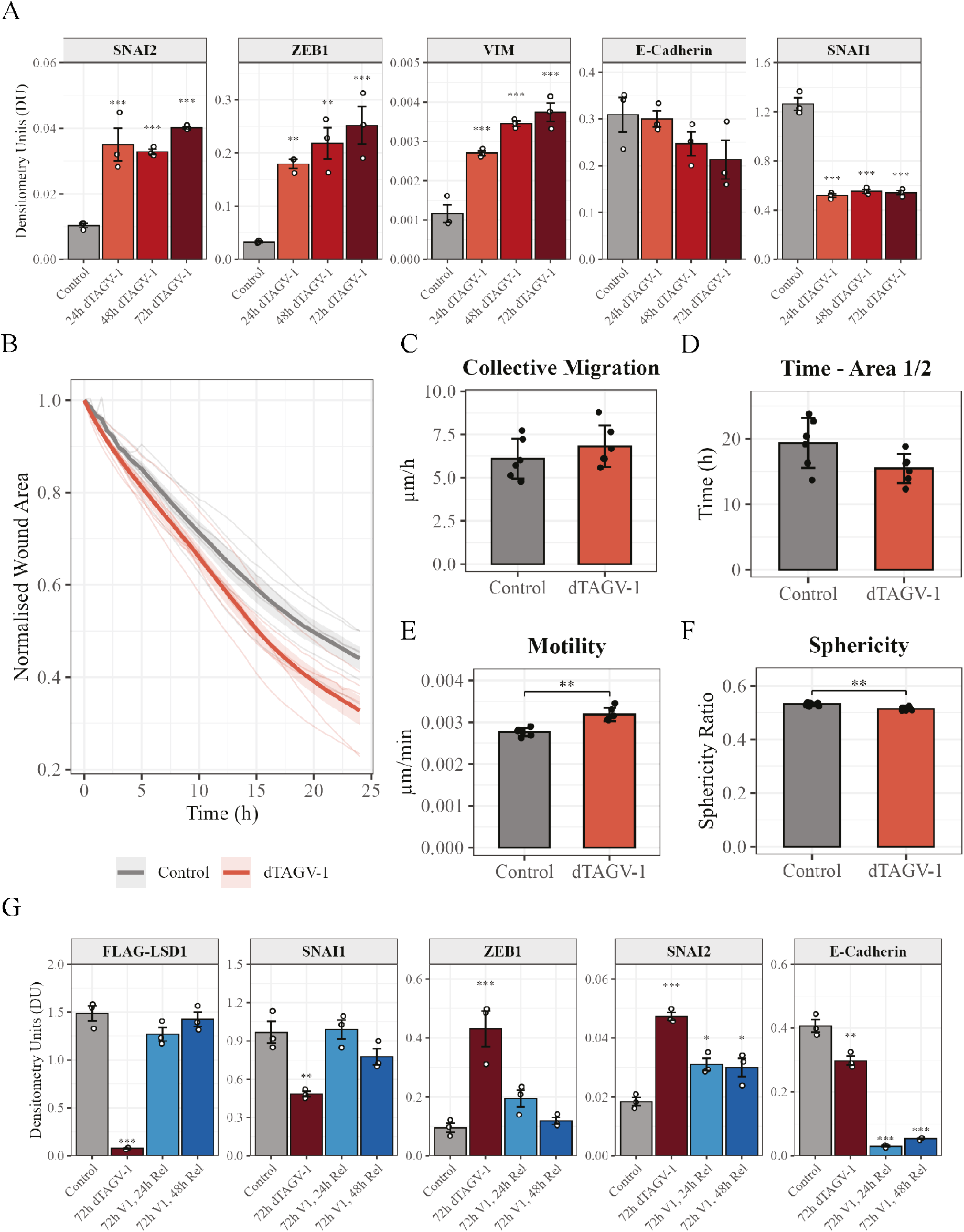
Depletion and re-introduction of LSD1 promotes a mesenchymal phenotype. (A) Densitometry of western blots for SNAI2, ZEB1, SNAI1 and E-cadherin following LSD1 depletion for 72 h; one-way ANOVA with post-hoc Dunnett test versus control, N = 3. (B) Normalised wound area over 24 h, (C) collective cell migration (µm/h), (D) time to 50% wound closure, (E) average cell motility (µm/min) and (F) average cell sphericity in control and 24 h dTAGV-1-treated cells, N = 6. (G) Densitometry of western blots for SNAI2, ZEB1, SNAI1, E-cadherin and FLAG following 72 h LSD1 depletion and 24 h and 48 h recovery; one-way ANOVA with post-hoc Tukey test versus control, N = 3. (A, G) TBP was used as a loading control. **p ≤ 0.01; ***p ≤ 0.001.

Consistent with a shift towards a partial mesenchymal identity, scratch-wound assays 24 h after LSD1 ablation showed faster wound closure, with increased collective cell migration and motility, reduced time to 50% wound closure and reduced cell sphericity (Fig. 4B–F).

We next asked whether this transition was stable or could be reverted by re-introducing LSD1, exploiting the swift restoration of protein levels following dTAG^V^-1 washout. After 72 h dTAG^V^-1 treatment, washout restored LSD1 and SNAI1 protein within 24 h, and ZEB1 fell to pre-treatment levels (Fig. 4G; Supplementary Fig. S5B). In contrast, SNAI2 decreased by ∼50% at 48 h but remained significantly above control (Fig. 4G; Supplementary Fig. S5B). Strikingly, E-cadherin did not recover but was further repressed (<10% of control) following LSD1 re-expression (Fig. 4G; Supplementary Fig. S5B).

Together this suggests LSD1 depletion induces EMT TFs and structural proteins, repressing E-cadherin and increasing migration, consistent with partial EMT. LSD1 re-introduction reverses the induction of mesenchymal genes within 48 h but further represses epithelial gene expression, possibly through LSD1 binding to residual SNAI2. In the tumour microenvironment, this could represent a further primed state able to undergo EMT in response to external factors.

### Distinct transcriptional signatures of LSD1 inhibition and LSD1 depletion

Finally, we asked whether LSD1 inhibition replicates LSD1 depletion. LSD1 also acts as a scaffold enabling CoREST HDAC1/2 deacetylation, so ablation could act through loss of demethylase and/or scaffolding and HDAC1/2 activity. To discriminate between these, we performed mRNA-seq on LSD1-FKBP12^F36V^-FLAG HCT116 cells 24 h after treatment with GSK-LSD1, which blocks LSD1 demethylase activity. PCA showed robust clustering of replicates with clear separation between DMSO- and GSK-LSD1-treated samples (Supplementary Fig. S6).

We expected demethylase inhibition to affect a subset of the genes responsive to LSD1 depletion, owing to retained HDAC activity. Indeed, although *ZEB1*, *VIM* and *SNAI2* were among the most upregulated genes after LSD1 depletion, *SNAI2* was unchanged following inhibition. LSD1 binds the promoters of all three genes (Fig. 5A; Supplementary Fig. S7A– C), but *SNAI2* also has LSD1 binding sites at upstream and downstream enhancers known to drive its transcription (Fig. 5A; Supplementary Fig. S7C) (Sundararajan et al. 2019; Pomella et al. 2021). Differential responses to depletion versus inhibition could therefore result from LSD1 remaining bound at regulatory elements, enabling repression via CoREST-associated HDAC1/2 activity.

**Figure 5.**
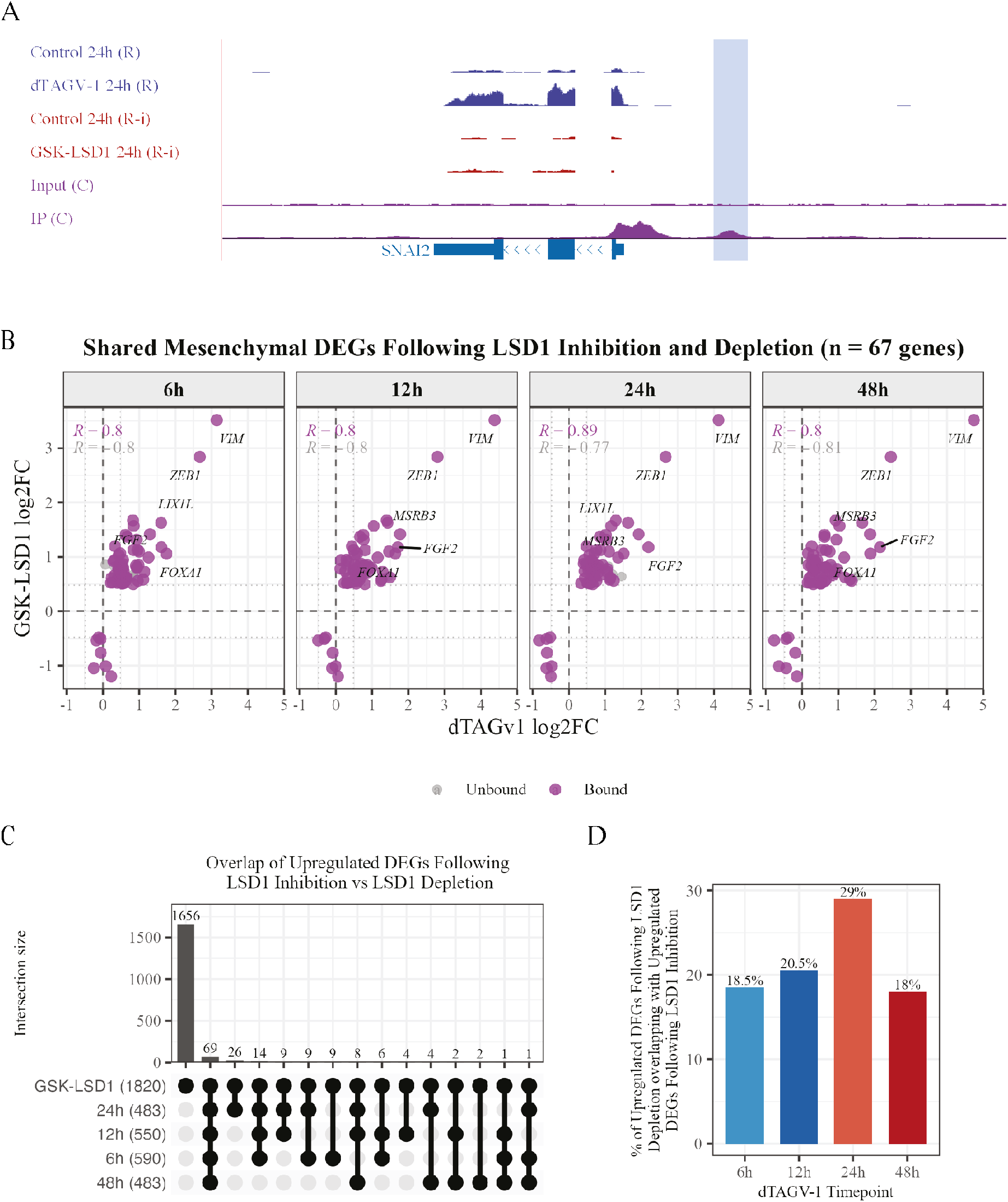
LSD1 inhibition produces a significantly different transcriptional response from LSD1 depletion. (A) Genome tracks across the SNAI2 gene following 24 h LSD1 depletion (dTAGV-1) or inhibition (GSK-LSD1), with LSD1 binding; an upstream enhancer element is highlighted in blue. (B) Bubble plot of log2 fold change of shared mesenchymal DEGs following LSD1 degradation at each timepoint (x-axis) and LSD1 inhibition (y-axis), with LSD1-bound genes in purple; the five genes with the largest log2 fold change are labelled. (C) UpSet plot of upregulated DEGs following LSD1 inhibition and LSD1 depletion, displayed as in Fig. 2A. (D) Bar plot of the percentage of upregulated DEGs following LSD1 depletion at each timepoint that are also upregulated following LSD1 inhibition.

At other loci, responses overlapped: fold changes of 67 mesenchymal mRNAs were highly correlated between treatments, with the largest changes in *ZEB1* and *VIM* (Fig. 5B). Surprisingly, however, GSK-LSD1 produced substantially more DEGs than LSD1 depletion: 1,820 upregulated and 1,686 downregulated mRNAs at 24 h (Supplementary Fig. S6B), compared with 483 upregulated and 54 downregulated 24 h after depletion. Consistent with this, there was limited overlap between DEGs following LSD1 inhibition and depletion (Fig. 5C-D and Supplementary Fig. S6C-D). Given the greater specificity of PROTAC-based targeting, the broader response to GSK-LSD1 may reflect, at least in part, off-target effects. However, more than 50% of DEGs following LSD1 inhibition overlapped LSD1-bound genomic regions (Supplementary Fig. S6E, F), suggesting that a substantial proportion of these changes may be associated with LSD1-bound regulatory regions. Pharmacological inhibition and targeted depletion of LSD1 therefore have distinct transcriptional consequences, underscoring the need to distinguish on-from off-target effects and to clarify the underlying mechanisms.

## Discussion

Stable switching of cell identity requires activation of the new lineage programme and silencing of the original one. Using rapid degron-mediated ablation of the CoREST subunit LSD1 with integrated multi-omics to characterise early, direct transcriptional and chromatin changes, we show that LSD1 has context-dependent roles in regulating transitions between epithelial and mesenchymal states. In addition to previously described roles in driving and maintaining mesenchymal phenotypes (Ferrari-Amorotti et al. 2013; Ferrari-Amorotti et al. 2014; Wang et al. 2015; Xie et al. 2020; Wang and Liu 2023), our data indicate that CoREST has a contradictory function in maintaining epithelial identity by repressing mesenchymal genes (Fig. 6).

**Figure 6.**
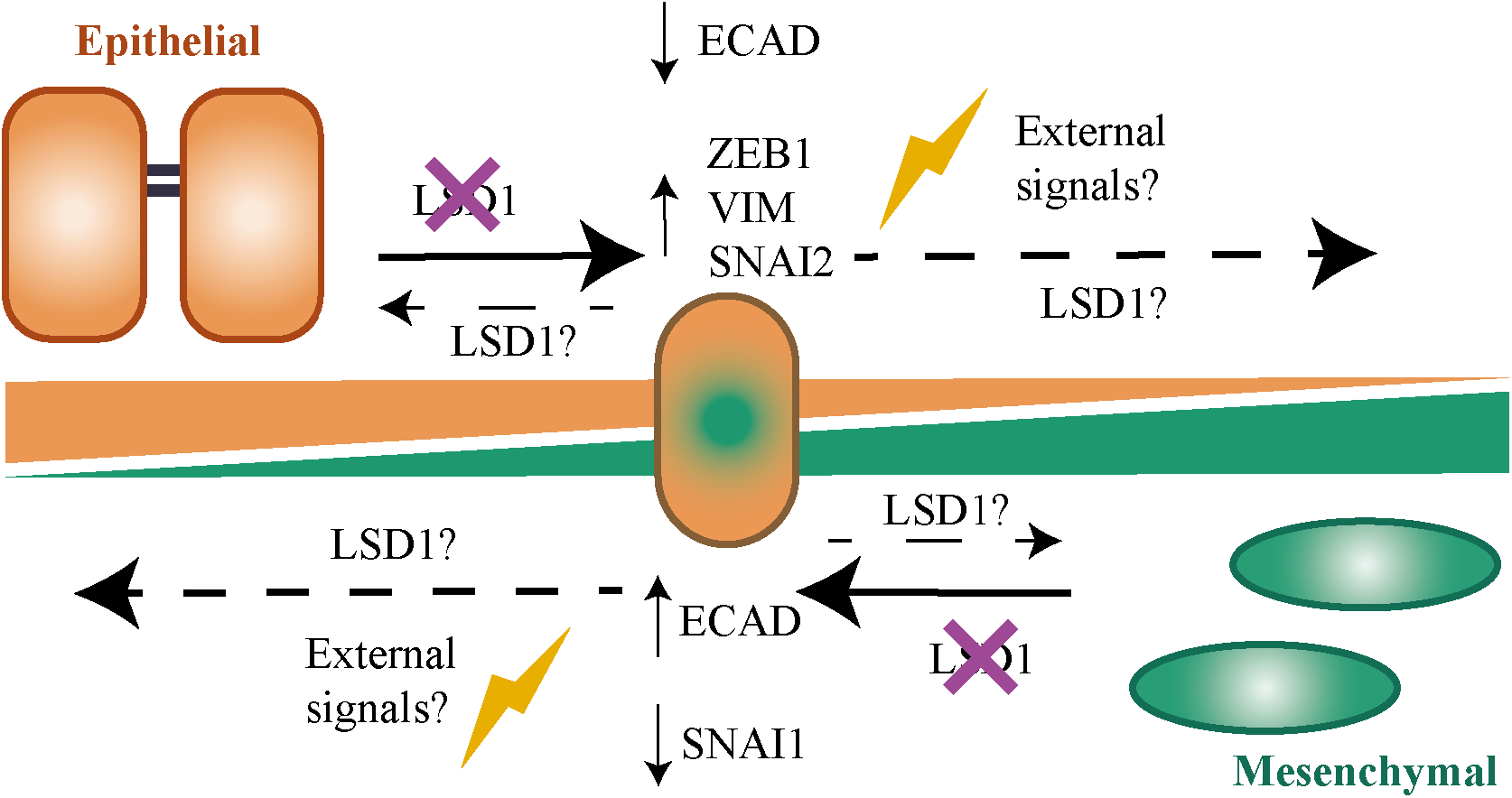
Schematic of the proposed role of LSD1 in maintaining epithelial and mesenchymal cell identity. Depletion or inhibition of LSD1 in either cell type results in partial EMT (epithelial cells) or partial MET (mesenchymal cells), producing an intermediate phenotype.

Although CoREST can repress genes through LSD1 demethylation of H3K4me1/2 and HDAC1/2 deacetylation of H3K9ac, or activate them through LSD1 demethylation of H3K9me1/2 (Shi et al. 2004; Lee et al. 2005; Metzger et al. 2005), the overwhelming majority of DEGs following LSD1 depletion were upregulated, confirming a direct repressive role; the increase in downregulated genes over time likely reflects secondary or non-specific effects. Approximately half of upregulated DEGs persisted from 6 h to 48 h, indicating rapid and sustained effects on transcription.

About one-third of upregulated genes were mesenchymal, more than 65% of which showed increased nascent transcription. In untreated cells, LSD1 localised to promoter-proximal regions within constitutively open chromatin, suggesting that LSD1 enforces a global programme of mesenchymal gene repression in epithelial cells. Strikingly, 60% of LSD1-bound genes and more than 90% of genes with increased steady-state mRNA were associated with constitutively active chromatin. LSD1/CoREST thus appears to bind open promoters of mesenchymal genes poised for rapid activation and restrain their transcription, at least in part, by reducing Pol II promoter occupancy. This is largely consistent with models in which co-repressors bind actively transcribed loci to maintain promoters in a state that, while not active, is primed for transcription (Reynolds et al. 2013).

Upregulated mesenchymal genes included *SNAI2*, *ZEB1* and *VIM*, which also increased at the protein level, while E-cadherin decreased, consistent with EMT activation and with previous reports of increased VIM and ZEB1, and decreased E-cadherin, following LSD1 inhibition (Choi et al. 2015; Yamakado et al. 2023). LSD1 bound the promoters of *SNAI2*, *ZEB1* and *VIM*, and additional LSD1 binding regions were observed at known enhancer elements upstream and downstream of *SNAI2* (Sundararajan et al. 2019; Pomella et al. 2021). LSD1 may therefore regulate genes through distinct mechanisms at promoters and enhancers, with CoREST-associated HDAC1/2 activity potentially particularly important at enhancers (Zeng et al. 2023).

Our data raise the possibility that LSD1 may execute contradictory roles in epithelial and mesenchymal states by being recruited to distinct promoters by non-overlapping TFs. SNAI1/2 and ZEB1/2 recruit LSD1 to epithelial genes in mesenchymal cells, but the factors recruiting LSD1 to mesenchymal genes in epithelial cells are less clear. In MDA-MB-231 mesenchymal cells, transient UTX expression has been suggested to recruit LSD1 to repress SNAI1, ZEB1 and ZEB2 (Choi et al. 2015), whereas in MCF-7 epithelial cells BRMS1 reportedly recruits LSD1 to repress VIM (Qiu et al. 2018). Although UTX binding in HeLa cells and LSD1 binding in HCT116 cells co-occurred at the *SNAI1* promoter, we detected no co-occupancy at *SNAI2*, *ZEB1* or *ZEB2*, and iRegulon did not identify UTX or BRMS1 motifs enriched in upregulated genes. However, ELF3 motifs were identified in genes upregulated at 6 h; ELF3 is reported to be a master epithelial regulator that represses EMT and antagonises *ZEB1* expression in HCT116 cells (Subbalakshmi et al. 2022). Different TFs may recruit LSD1 in epithelial and mesenchymal cells. However, SNAI1, SNAI2 and ZEB1 motifs were also enriched in upregulated genes and SNAI2 and ZEB1 proteins increased after LSD1 depletion, suggesting an alternative feed-forward mechanism in which LSD1 represses mesenchymal genes whose activation also relies on SNAI2 and ZEB1 upregulation.

An advantage of the dTAG^V^-1 system is that it enables the swift restoration of protein levels following dTAG^V^-1 washout, allowing us to test the stability of transcriptome and phenotype switches that occur following LSD1 depletion. E-cadherin fell after LSD1 depletion and, rather than returning to control levels, was further repressed following LSD1 re-introduction, suggesting that epithelial cells partially enter EMT. We speculate that the initial partial repression of E-cadherin in the absence of LSD1 could involve additional TFs such as TCF12 (Lee et al. 2012), whose motifs were enriched in upregulated genes at all timepoints. When LSD1 is restored, E-cadherin is further repressed through an LSD1–SNAI1/SNAI2-dependent mechanism, advancing the transition towards EMT.

LSD1 depletion and re-expression may therefore push cells towards a mesenchymal phenotype, providing a window of ≥48 h during which full EMT could occur in the presence of external factors. Whether this applies to LSD1 inhibition remains to be determined, but it suggests caution should be taken when considering LSD1 inhibition therapeutically. Short-term inhibition is likely to reduce migration and invasion of cells that already display a mesenchymal phenotype, thereby reducing metastatic potential. However, epithelial cells within a tumour microenvironment may undergo a partial EMT that progresses to full EMT when inhibition is removed. Consistent with this, whereas LSD1 inhibition in metastatic cancer cell lines reduces migration and invasion (Ferrari-Amorotti et al. 2013; Ferrari-Amorotti et al. 2014; Wang et al. 2015; Xie et al. 2020; Wang and Liu 2023), we observed small increases in cell migration and motility and reduced cell sphericity, in accordance with oral squamous carcinoma cell lines in which chemical inhibition of LSD1 triggers full EMT with exogenous SNAI1 expression (Yamakado et al. 2023).

A further note of caution regarding therapeutic LSD1 inhibitors is that fewer than 6% of upregulated and less than 1% of downregulated genes following LSD1 inhibition overlapped with those observed after LSD1 depletion, although 63% of upregulated and 56% of downregulated genes following inhibition exhibited LSD1 binding. GSK-LSD1 has been reported to inhibit both LSD1 activity and chromatin binding (Maiques-Diaz et al. 2018), yet approximately 50% of LSD1 binding sites remain occupied after inhibitor treatment (Egolf et al. 2019). Chromatin-bound, inhibited LSD1 may therefore preserve CoREST-associated HDAC activity, in agreement with previous studies (Song et al. 2020), or perform an enzymatically independent role (Zeng et al. 2023).

These results add another layer to the complex role LSD1 plays within epithelial and mesenchymal phenotypes. While inhibiting LSD1 in mesenchymal cells is widely accepted to partially reverse EMT, our data suggest that LSD1 maintains both epithelial and mesenchymal identities by globally repressing genes associated with the alternative phenotype, similar to its role in repressing neuronal genes in non-neuronal cells (Ballas et al. 2001). When LSD1 is inhibited in either cell type, cells may revert to an intermediate state expressing both epithelial and mesenchymal markers. In cell culture, these changes are reversible, but in disease, the complex tumour microenvironment could influence progression towards EMT or MET (Fig. 6). Addressing these questions is essential to understand the short- and long-term effects of targeting LSD1 in cancer.

## Methods

### Cell culture and proliferation

Parental HCT116 and CRISPR/Cas9-engineered HCT116.LSD1FK/FL cells were cultured for a maximum of four weeks in Dulbecco’s modified Eagle’s medium (DMEM; ThermoFisher) supplemented with 10% foetal bovine serum (Sigma) and 1% penicillin, streptomycin, glutamine (PSG; Sigma), and detached using TrypLE (Invitrogen™). Cells were treated for 2-72 h with 100 nM dTAG^V^-1 (R&D Systems), 100 nM GSK-LSD1 (R&D Systems) or an equivalent volume of DMSO. To measure proliferation, 0.15 x 10^6^ cells were seeded into 6-well plates and counted every 24 h using a BioRad automated cell counter.

### Preparation of samples for western blotting

Whole-cell extracts were prepared by resuspending cell pellets in RIPA buffer (50 mM Tris/Cl pH 7.4, 150 mM NaCl, 1% IGEPAL CA-630, 0.5% sodium deoxycholate, 0.1% SDS, 1 mM EDTA), rotating end-over-end at 4 °C for 30 min and removing insoluble material by centrifugation (15,000 RCF, 20 min, 4 °C). To isolate soluble nuclear proteins, cell pellets were resuspended in lysis buffer 1 (10 mM Tris/Cl pH 7.4, 1 mM MgCl2, 0.1% IGEPAL CA-630, 0.5 mM DTT), vortexed and incubated on ice for 10 min. Nuclei were collected by centrifugation (1,000 RCF, 3 min, 4 °C), resuspended in lysis buffer 2 (lysis buffer 1 with 500 mM NaCl) and incubated on ice for 15 min with vortexing every 5 min. Precipitated DNA and insoluble material were removed by centrifugation (5,000 RCF, 3 min, 4 °C). All buffers were supplemented with EDTA-free protease inhibitor cocktail (Roche). Protein concentration was determined using BioRad protein assay reagent.

### Western blotting

Similar amounts of protein were run on NuPAGE 4–12% Bis-Tris gels in 1x MES SDS running buffer or 4–8% Tris-acetate gels in 1x Tris-acetate running buffer (Invitrogen™) and wet-transferred to nitrocellulose membranes in transfer buffer (Invitrogen™) with 10–20% methanol at 30 mA for 1 h. Tris-acetate gels were pre-incubated in 2x transfer buffer with 0.02% SDS for 10 min and transferred for 2 h with 0.01% SDS in the transfer buffer.

Membranes were air-dried for 30 min, blocked for 1 h in Tris-buffered saline (TBS; 20 mM Tris, 150 mM NaCl, pH 7.6) with 5% milk, and incubated overnight at 4 °C with primary antibodies (Ms CoREST, 1:500, Millipore; Rb E-cadherin, 1:1,000, Cell Signalling; Ms FLAG, 1:1,000, Sigma; Rb HDAC1, 1:1,000, Abcam; Rb LSD1, 1:1,000, Abcam; Rb SNAI1, 1:1,000, Cell Signalling; Rb SNAI2, 1:1,000, Cell Signalling; Rb ZEB1, 1:1,000, Abcam; Ms TBP, 1:300, Santa Cruz) and for 1 h at room temperature with secondary antibodies (IRDye 680RD Gt anti-mouse or IRDye 800CW Gt anti-rabbit, both 1:20,000; LI-COR). Membranes were washed 3 x 10 min with TBS containing 0.1% Tween-20 before antibody incubations and imaging, imaged on an Odyssey CLx and processed using Image Studio (v5.5.4).

### Preparation of samples for RNA sequencing

Cells were seeded into 6-well plates and allowed to settle for 24 h before treatment with dTAG^V^-1 for 6–48 h. Cells were washed with PBS, detached using TrypLE, diluted in PBS, transferred to nuclease-free 1.5 ml tubes and collected by centrifugation (400 RCF, 5 min, room temperature). Total RNA was isolated using a Maxwell RSC (Promega) and quality-checked by the Nucleus Genomics facility, University of Leicester. Each experiment was carried out in triplicate.

### Processing RNA-sequencing data

RNA sequencing was performed by Novogene on a NovaSeq 6000 S4 platform (150 bp paired-end, depth 20 million reads). Files were quality-checked using FastQC ((Andrews 2010), v0.12.1) and aligned using HISAT2 ((Kim et al. 2019a), v12.3.0) with standard options to the human genome build GRCh38 release 111 (Dyer et al. 2025). Alignments were converted to BAM format, coordinate-sorted and indexed using Samtools ((Li et al. 2009), v1.17). Count files were generated using LiBiNorm ((Dyer et al. 2019), v2.5; options: -z -r pos -I gene_name -s no) and the GRCh38 release 111 gene transfer format file.

Unless stated otherwise, analysis was performed in Rstudio using R (v4.4.1). Differential expression was performed with DESeq2 ((Love et al. 2014), v1.44.0; Wald test, local fit type, post-hoc Benjamini-Hochberg P-value correction), with apeglm ((Zhu et al. 2019); v1.26.1) providing shrinkage estimates of log fold changes. Genes with counts less than 50 in 3 or more samples were excluded. Gene set enrichment analysis (GSEA) (Mootha et al. 2003; Subramanian et al. 2005) was tested against the MSigDB hallmark gene sets. Over-represented TF binding motifs within 500bp of the TSS were identified with iRegulon ((Janky et al. 2014), v1.3) with standard options and used to create a network map in cytoscape ((Shannon et al. 2003), v3.10.5).

### Preparation of samples for quick precision run-on (qPRO) sequencing

Samples were prepared for qPRO-seq as previously described (Judd et al. 2020) (dx.doi.org/10.17504/protocols.io.57dg9i6). Treated cells were washed twice with ice-cold PBS and scraped into ice-cold permeabilisation buffer (10 mM Tris/Cl pH 8, 10 mM KCl, 250 mM sucrose, 5 mM MgCl2, 1 mM EGTA, 0.1% IGEPAL CA-630, 0.5 mM DTT, 0.05% Tween-20, 10% glycerol, PIC and SUPERase-In RNase inhibitor (ThermoFisher)). Cells were incubated on ice for 5 min, collected by centrifugation (700 RCF, 4 min, 4 °C), washed twice in permeabilisation buffer without IGEPAL CA-630 and Tween-20, resuspended in freeze buffer (50 mM Tris/Cl pH 8, 40% glycerol, 5 mM MgCl2, 1.1 mM EDTA, 0.5 mM DTT, SUPERase-In RNase inhibitor) at 1 x 10^6^ cells/52 µl and snap-frozen in aliquots. For the run-on reaction, 50 µl permeabilised cells were mixed 1:1 with 2x run-on master mix (10 mM Tris/Cl pH 8, 5 mM MgCl2, 1 mM DTT, 300 mM KCl, 40 µM each of Biotin-11-CTP/UTP/ATP/GTP (PerkinElmer), 1% sarkosyl, SUPERase-In RNase inhibitor) and incubated at 37 °C, 750 rpm for exactly 5 min. The reaction was stopped with RL buffer and RNA purified following the manufacturer’s instructions (Norgen Biotek Corp.). Purified RNA was hydrolysed with 1 M NaOH, ethanol-precipitated and resuspended in nuclease-free water. 3′ RNA adaptors were ligated using T4 RNA ligase 1 (NEB) and biotin-labelled RNA bound to streptavidin beads (Invitrogen™), followed by on-bead 5′ hydroxyl repair with T4 polynucleotide kinase (NEB), 5′ decapping with RppH (NEB) and 5′ RNA adaptor ligation with T4 RNA ligase 1. RNA was eluted with TRIzol and reverse-transcribed off-bead using Maxima H Minus RT (ThermoFisher). Following test amplification, full-scale amplification was performed using Q5 high-fidelity DNA polymerase (NEB). DNA was purified using AMPure beads (Invitrogen™), quantified using the Qubit HS dsDNA assay (Invitrogen™) and quality-checked on an Agilent 2100 Bioanalyzer (Agilent Technologies). RNase-free water, LoBind RNase-free tubes and wide-bore nuclease-free filter tips were used throughout.

### Processing qPRO-sequencing data

qPRO sequencing was performed by Novogene on a NovaSeq 6000 S4 (150 bp paired-end, minimum depth 50 million reads) and files were quality-checked using FastQC. Fastp ((Chen et al. 2018), v0.23.2) was used for preprocessing and Illumina adaptor removal (options: --umi --stdout --umi_loc=per_read --umi_len=6 -c --overlap_len_require 15) and bowtie2 ((Langmead and Salzberg 2012), v12.3.0) to remove reads aligning to ribosomal RNA (options: --fast-local --un-conc --interleaved). Reads were aligned to the human hg38 genome using bowtie2 (options: --local --sensitive-local), reads with MAPQ <10 removed, and files converted to BAM format using Samtools. BAM files were deduplicated using UMI-tools ((Smith et al. 2017), v1.0.1; options: --umi-separator=“:” --paired) then sorted and indexed using Samtools. BigWig files of forward and reverse reads were created using deepTools ((Ramirez et al. 2014), v3.5.1) bamCoverage (options: --skipNonCoveredRegions --binSize 1 --normalizeUsing None --Offset 1 --samFlagInclude 82 (forward) or 98 (reverse)).

BigWig files were imported into RStudio using BRGenomics ((DeBerardine 2023), v1.14.1). Gene boundaries were generated from TxDb.Hsapiens.UCSC.hg38.knownGene, with gene bodies defined from TSS +300 bp to transcription end site (TES) −300 bp, and promoters as TSS ±500 bp. BRGenomics was used to subsample for metaplots and create count matrices. Differential expression was performed using DESeq2 with apeglm shrinkage as for RNA-seq, excluding genes with counts less than 20 in 2 or more samples.

### Preparation of samples for ATAC sequencing

Following treatment, cells were detached and counted, and 100,000 cells per condition were washed once in cold PBS. ATAC-seq libraries were prepared using the Active Motif ATAC-seq kit. Briefly, cells were lysed and tagmentation performed at 37 °C, 800 rpm for 30 min. Tagmented DNA was column-purified, PCR-amplified using i7 and i5 indexed primers and cleaned up using SPRI beads, and size distribution was assessed on an Agilent 2100 Bioanalyzer.

### Processing ATAC-sequencing data

ATAC sequencing was performed by Novogene on a NovaSeq 6000 S4 (150 bp paired-end, minimum depth 50 million reads) and files were quality-checked using FastQC. Adaptors were trimmed with TrimGalore (v0.6.6; options: --paired) and reads aligned to hg38 using bowtie2 (options: --very-sensitive -X 1000). Unknown and mitochondrial reads were removed, and Samtools was used to convert to BAM format and restrict to properly paired reads (Li et al. 2009). Sorted, indexed datasets were subsampled with Samtools based on library complexity calculated using ATACseqQC ((Ou et al. 2018), v1.26), then sorted and indexed, and duplicates removed using Picard (v2.6.0) MarkDuplicates (options: REMOVE_DUPLICATES=true). Files were sorted by read name, missing paired-end information filled in using Samtools, and converted to BEDPE format using BEDTools ((Quinlan and Hall 2010) v2.30.0). Peaks were called with MACS2 ((Zhang et al. 2008), v2.2.6) callpeak (options: -f BEDPE -g hs --broad --broad-cutoff 0.05 --keep-dup all) and filtered to exclude blacklisted regions using BEDTools. BroadPeak, BAM and blacklist files were imported into RStudio, and significant differential chromatin accessibility was calculated using csaw ((Lun and Smyth 2016), v1.36.1) and edgeR ((Robinson et al. 2010), v4.0.16).

### Additional R packages

The following R packages were used to analyse and visualise the data: GenomicRanges (v1.62.1) (Lawrence et al. 2013); ChIPseeker (1.46.1) (Yu et al. 2015); ChIPpeakAnnno (v3.44.0) (Zhu et al. 2010); biomaRt (v2.66.2) (Durinck et al. 2005); org.Hs.eg.db (v3.22.0) (Carlson 2025); ggh4x (v0.3.1) (van den Brand 2024); ComplexHeatmap (v2.26.1) (Gu et al. 2016); VennDiagram (v1.8.2) (Chen and Boutros 2011); ggplot2 (v4.0.3) (Wickham 2009); ggupset (v0.4.1) (Ahlmann-Eltze 2025); ggcorrplot (v0.1.4.1) (Kassambara 2023); clusterProfiler (v4.18.4) (Wu et al. 2021)

### Statistics

Data are mean ± SD; significance was tested by one-way ANOVA with post-hoc Tukey or Dunnett tests for multiple comparisons using the R packages stats (v4.3.3) and multcomp (v1.4-25).

## Data Availability

The RNA-seq, qPRO-seq and ATAC-seq datasets produced in this study are available at Gene Expression Omnibus and will be provided upon publication.

LSD1 ChIP-seq data were previously published as GSE330806 (files GSM9733045– GSM9733048) (https://www.ncbi.nlm.nih.gov/geo/query/acc.cgi?acc=GSE330806).

## Acknowledgements

This work was supported by funding from the following sources: MRC studentship [to K.S.]; BBSRC Midlands Integrative Biosciences Training Partnership [to M.E.C.]; BBSRC [BB/L00996X/1; BB/M028054/1 to S.K. and P.B.]; Leverhulme Trust [RPG 2020-099 to S.K. and P.B.]; MRC [MR/Z506059 to J.T.H]; Wellcome Trust Investigator Award [222493/Z/21/Z to J.W.R.S.]; MRC [MR/W00190X/1 to S.M.C.]. We thank Nicolas Sylvius from the Nucleus Genomics facility at the University of Leicester for extracting RNA for sequencing and Louise Fairall for comments on the manuscript.

## Author contributions

R.E.T. Performed the RNA-seq and qPRO-seq dTAG^V^-1 experiments, processed RNA-seq, qPRO-seq, and ATAC-seq data, performed the western blotting, designed the figures and wrote the manuscript; O.G. Generated the LSD1-targeted HCT116 cell line; K.S. Performed the ATAC-seq experiments; M.I. performed the western blotting; M.E.C. performed the RNA-seq GSK-LSD1 inhibitor experiments; S.K. Performed the qPRO-seq experiments; J.T.H. Designed the inhibitor experiments and provided supervision; P.B. designed the qPRO-seq experiments and wrote the manuscript; S.M.C. designed the inhibitor experiments and provided project guidance; J.W.R.S. designed the experiments, provided supervision and wrote the manuscript.

