## Supplementary figures for "Global repression of mesenchymal genes in epithelial cells by the histone demethylase LSD1"

Robert E. Turnbull[1,2*], Oksana Gonchar[1,2], Kristupas Sirvydis[1,2], Mariam Iftikhar[1,2], Megan E. Coulson[2], So Yeon Kwon[3], James T. Hodgkinson[1,4], Paul Badenhorst[3], Shaun M. Cowley[2], John W.R. Schwabe[1,2*]

[1]Institute for Structural and Chemical Biology, [2]Division of Molecular and Cell Biology and [4]School of Chemical, Earth and Environmental Sciences, University of Leicester, Leicester, LE1 7RH, UK

[3]Institute of Cancer and Genomic Sciences, University of Birmingham, Birmingham, B15 2TT, UK

*Co-corresponding authors

**ORCID IDs:**

| | |
|---|---|
| Turnbull: | 0000-0003-2863-4374 |
| Gonchar: | 0000-0002-7835-5712 |
| Sirvydis: | 0000-0002-6337-3504 |
| Iftikhar: | 0009-0009-5967-5292 |
| Coulson: | 0000-0003-1058-4241 |
| Kwon: | 0000-0002-8490-9101 |
| Hodgkinson: | 0000-0001-9978-7322 |
| Badenhorst: | 0000-0002-2542-0250 |
| Cowley: | 0000-0002-2510-1305 |
| Schwabe: | 0000-0003-2865-4383 |

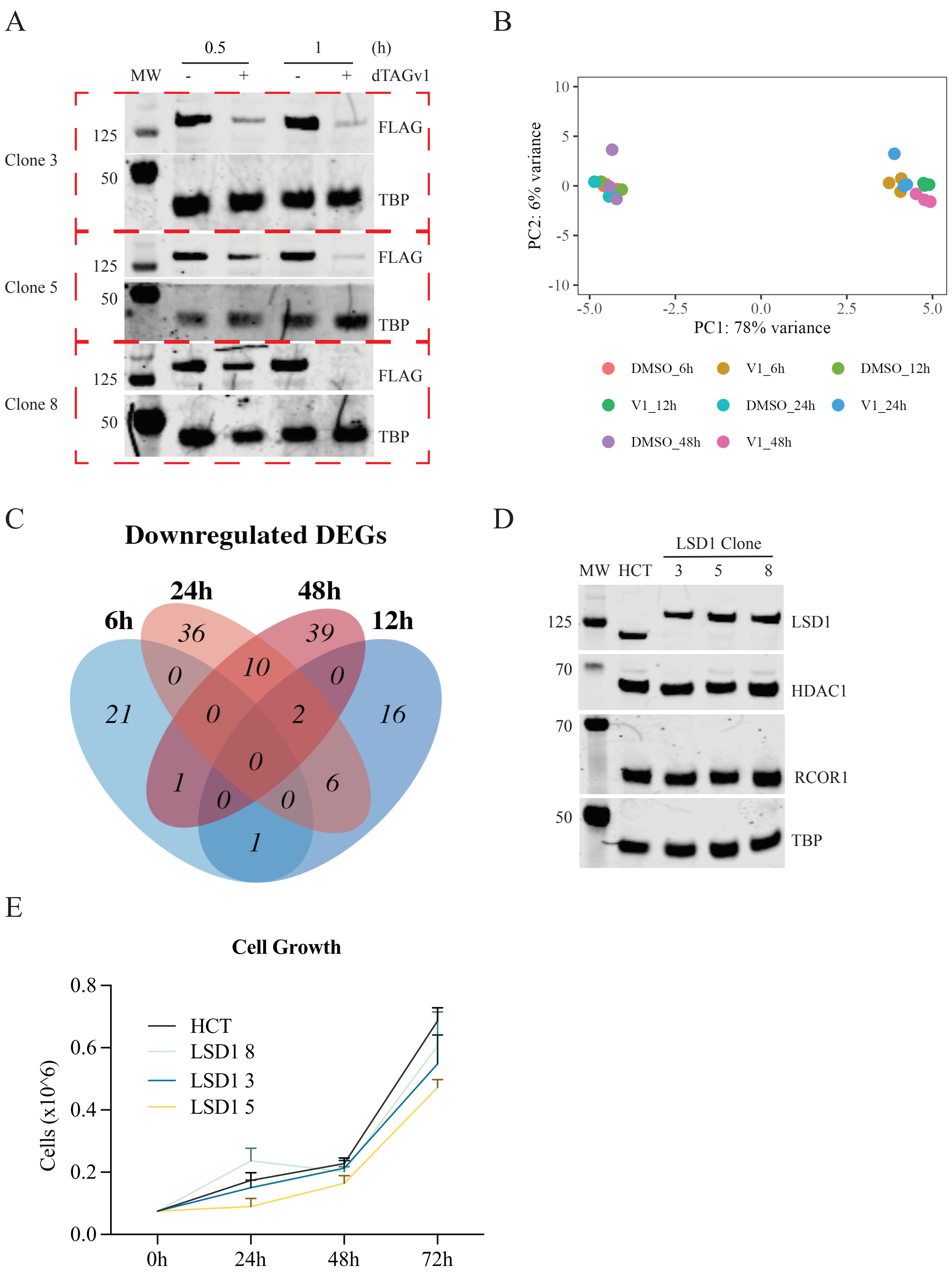


**Supplementary Fig. 1** CRISPR targeting of endogenous LSD1 enables rapid PROTAC-mediated degradation. (A) Western blot for FLAG following 30min or 1h treatment with PROTAC dTAG$^{V}$-1 of 3 independently derived HCT116 clones with tagged LSD1. DMSO was used as a vehicle control. TBP was used as a loading control. (B) Principle component analysis following RNA-sequencing of 6-48h dTAG$^{V}$-1-treated LSD1-targeted cells (Clone 3). (C) Venn diagram of downregulated differentially expressed genes following LSD1-depletion over 48h. (D) Western blot for LSD1, HDAC1 and RCOR1 in parental HCT116 and 3 independently-derived LSD1-targeted cell lines. TBP was used as a loading control. (E) Cell growth of parental and 3 independently-derived LSD1-targeted cell lines over 72h without treatment. n=3. [dTAG$^{V}$-1] = 100nM.

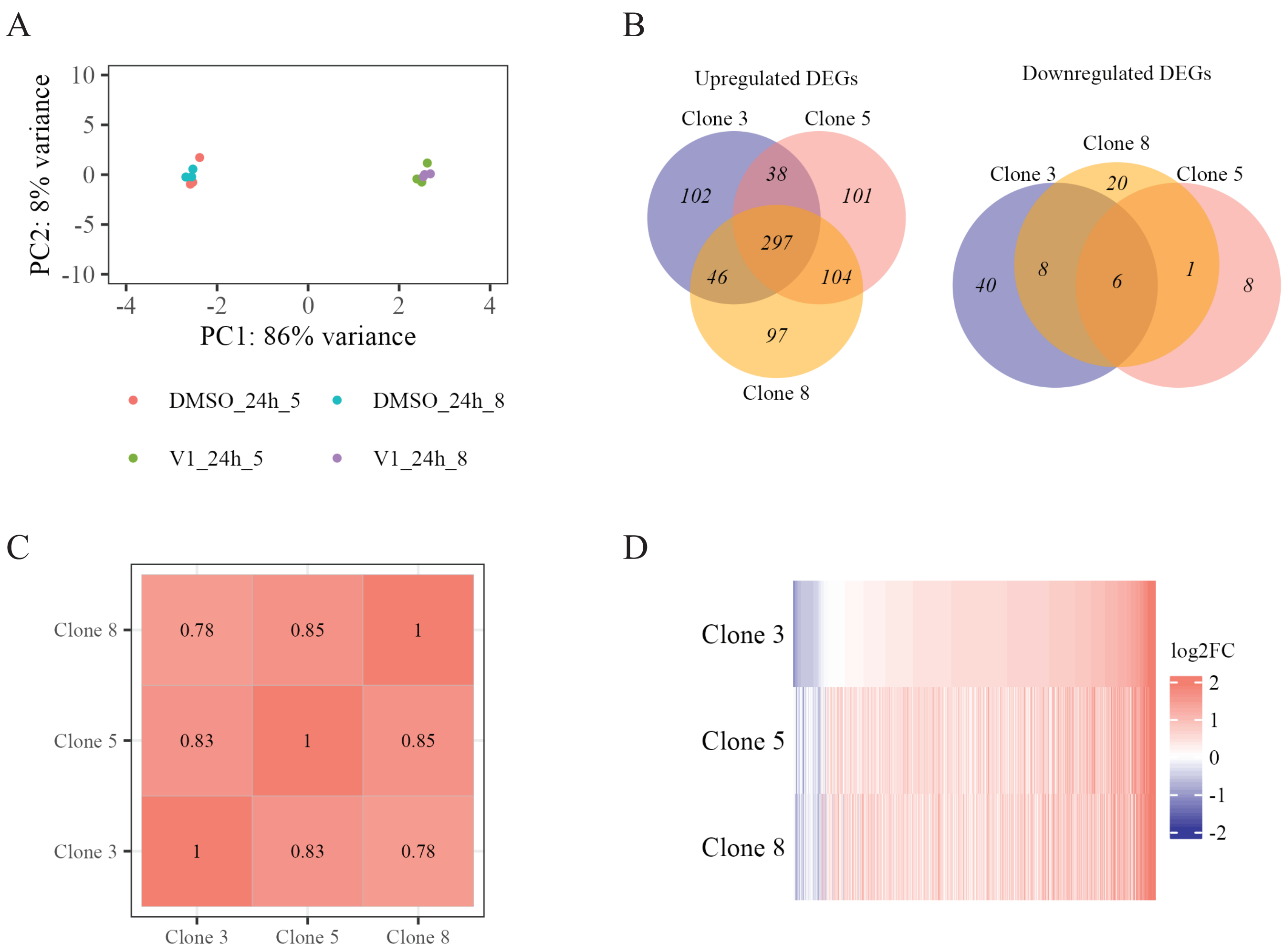


**Supplementary Fig. 2** Independent LSD1-targeted HCT116 clones exhibit similar changes in upregulated differentially expressed genes. (A) Principal component analysis of a further two independent clones following 24h dTAG$^{V}$-1 treatment. (B) Venn diagram of up and down differentially expressed genes from three independent clones following 24h dTAG$^{V}$-1 treatment. (C) Correlation matrix of all differentally expressed genes from all three independent clones following 24h dTAG$^{V}$-1 treatment. (D) Heatmap of log2 fold change of all differentially expressed genes from all three independent clones. [dTAG$^{V}$-1] = 100nM

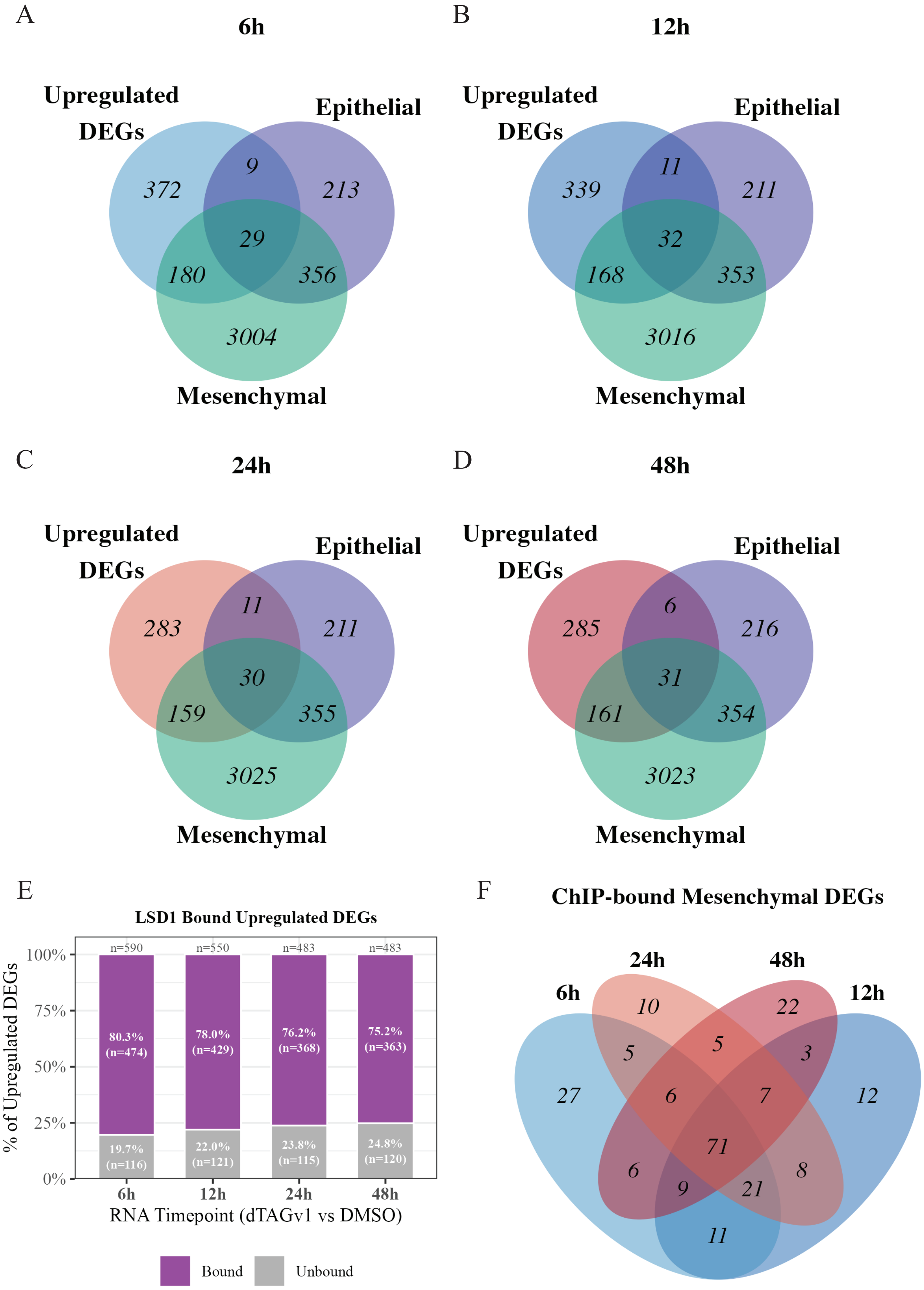


**Supplementary Fig. 3** Upregulated genes following LSD1 depletion are associated with a mesenchymal signature. (A-D) Venn diagrams of upregulated differentially expressed genes at each timepoint intersected with epithelial and mesenchymal genes curated from the EMTome. (E) Bar plot showing upregulated differentially expressed genes at each timepoint that have LSD1 bound. (F) Venn diagram of mesenchymal upregulated differentially expressed genes following LSD1 depletion at each timepoint that have bound LSD1.

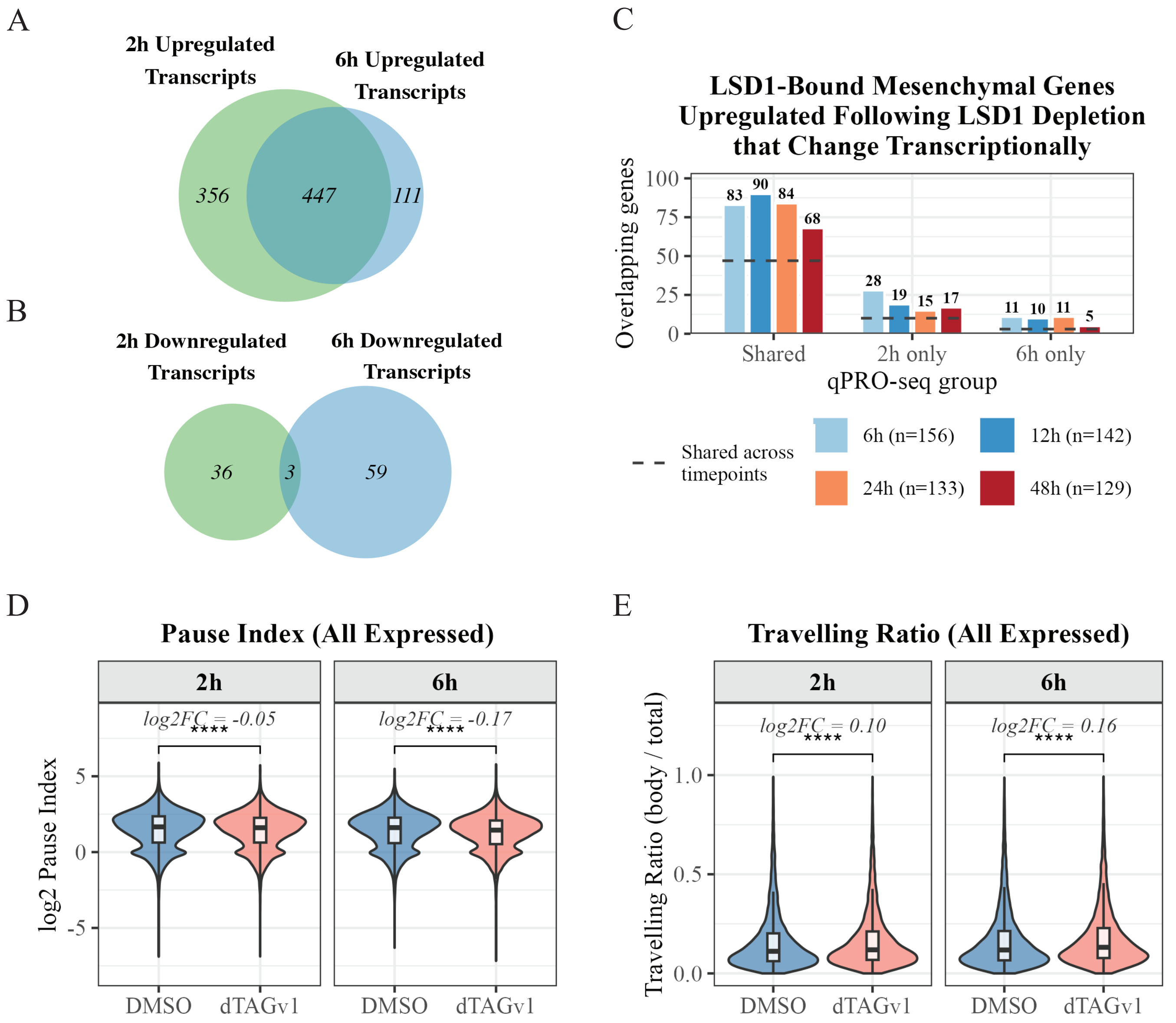


**Supplementary Fig. 4** LSD1 depletion leads to rapid increases in gene transcription. (A-B) Venn diagrams of intersecting genes that increase (A) or decrease (B) transcriptionally at 2h and 6h. (C) Bar plot showing the LSD1-bound mesenchymal upregulated differentially expressed mRNA genes at each timepoint following LSD1 degradation that show changes in transcription at both 2h and 6h, only 2h or only 6h. Horizontal dashed lines indicate the number of genes shared across all timepoints. (D) Pause index of all genes transcribed at 2h and 6h in the control versus LSD1 depletion. Thisis calculated as the number of reads ±300bp of the gene transcription start site divided by the number of reads over the gene body (+300bp from TSS, -300bp from TES). (E) Travelling ratio of all genes transcribed in 2h and 6h control versus LSD1 depletion. This is calculated as the number of reads over the gene body divided by the total number of reads. ****$p \leq 0.0001$.

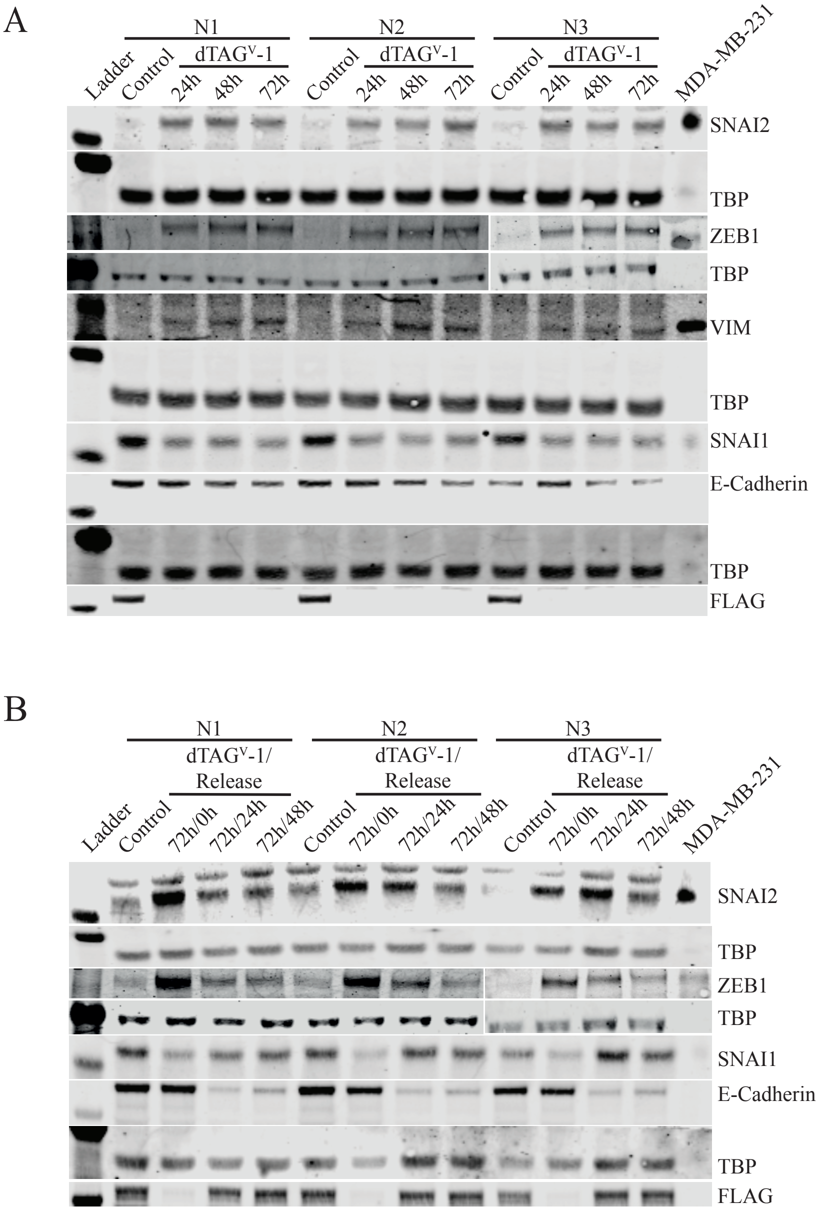


**Supplementary Fig. 5** LSD1 depletion and re-introduction drives a mesenchymal phenotype. (A) Western blot for SNAI2, ZEB1, VIM, SNAI1, E-Cadherin and FLAG following 24h, 48h, and 72h LSD1 depletion. (B) Western blot for SNAI2, ZEB1, SNAI1, E-cadherin and FLAG following 72h LSD1 depletion followed by a 24h or 48h release from dTAG$^{V}$-1 (through washing). MDA-MB-231 lysates were used as a postivie control, TBP was used as a loading control. N=3.

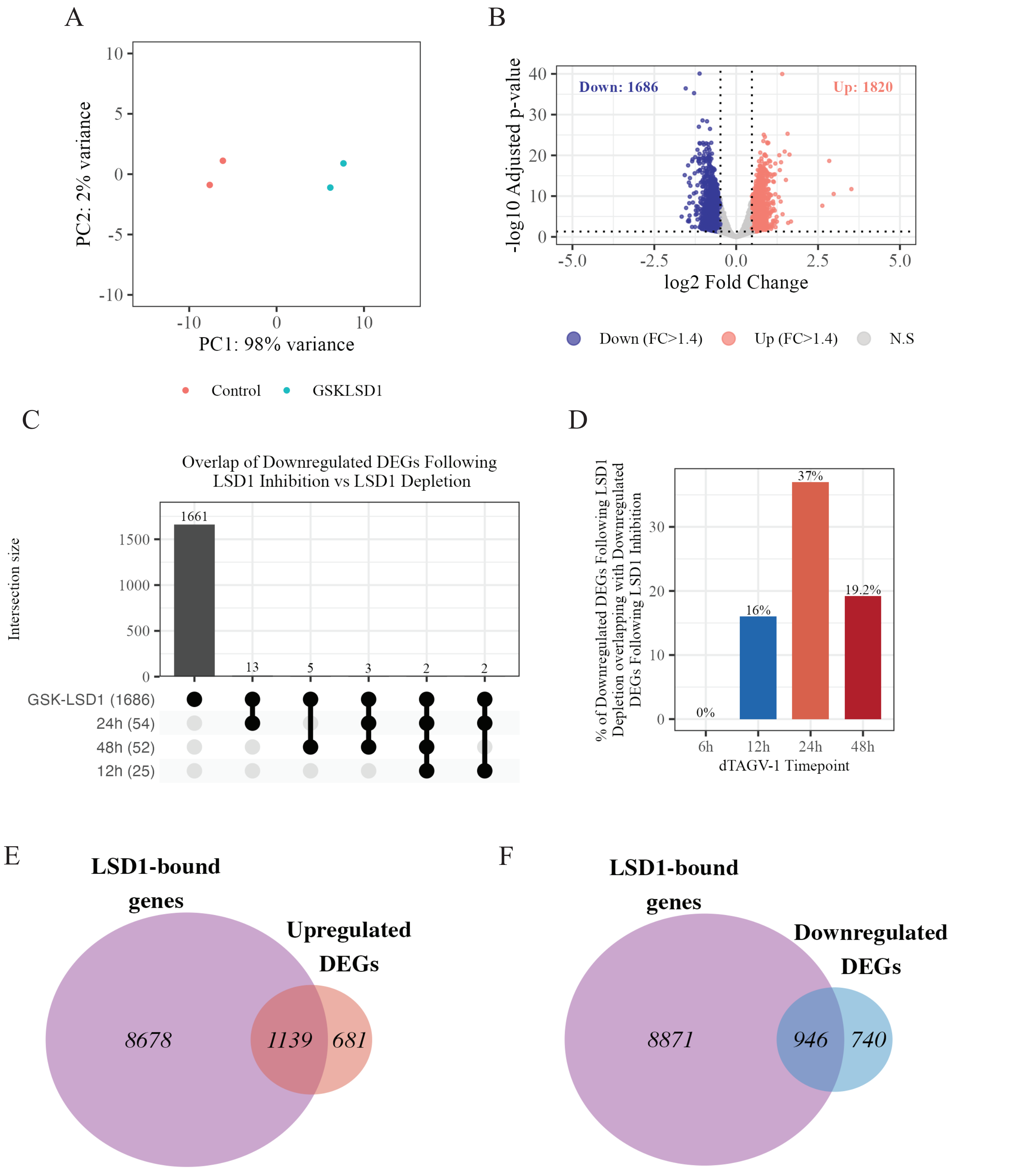


**Supplementary Fig. 6** LSD1 inhibition shows a distinct transcriptional signature compared to LSD1 depletion. (A) Principal component analysis showing the variation between duplicate control and 24h GSK-LSD1-treated samples using LSD1-targeted HCT116 cells. (B) Volcano plot showing the down (blue) and up (red) differentially expressed genes following LSD1 inhibition (pAdj $\leq$0.05, log2FC $\geq$1.4). (C) Upset plot of downregulated differentially expressed genes following LSD1 inhibition and and LSD1 depletion. (D) Bar plot showing percent of downregulated differentially expressed genes at each timepoint following LSD1 depletion that are downregulated following LSD1 inhibition. (E-F) Venn Diagrams of upregulated (E) and downregulated (F) differentially expressed genes following LSD1 inhibition overlapped with genes that show LSD1 genomic binding.

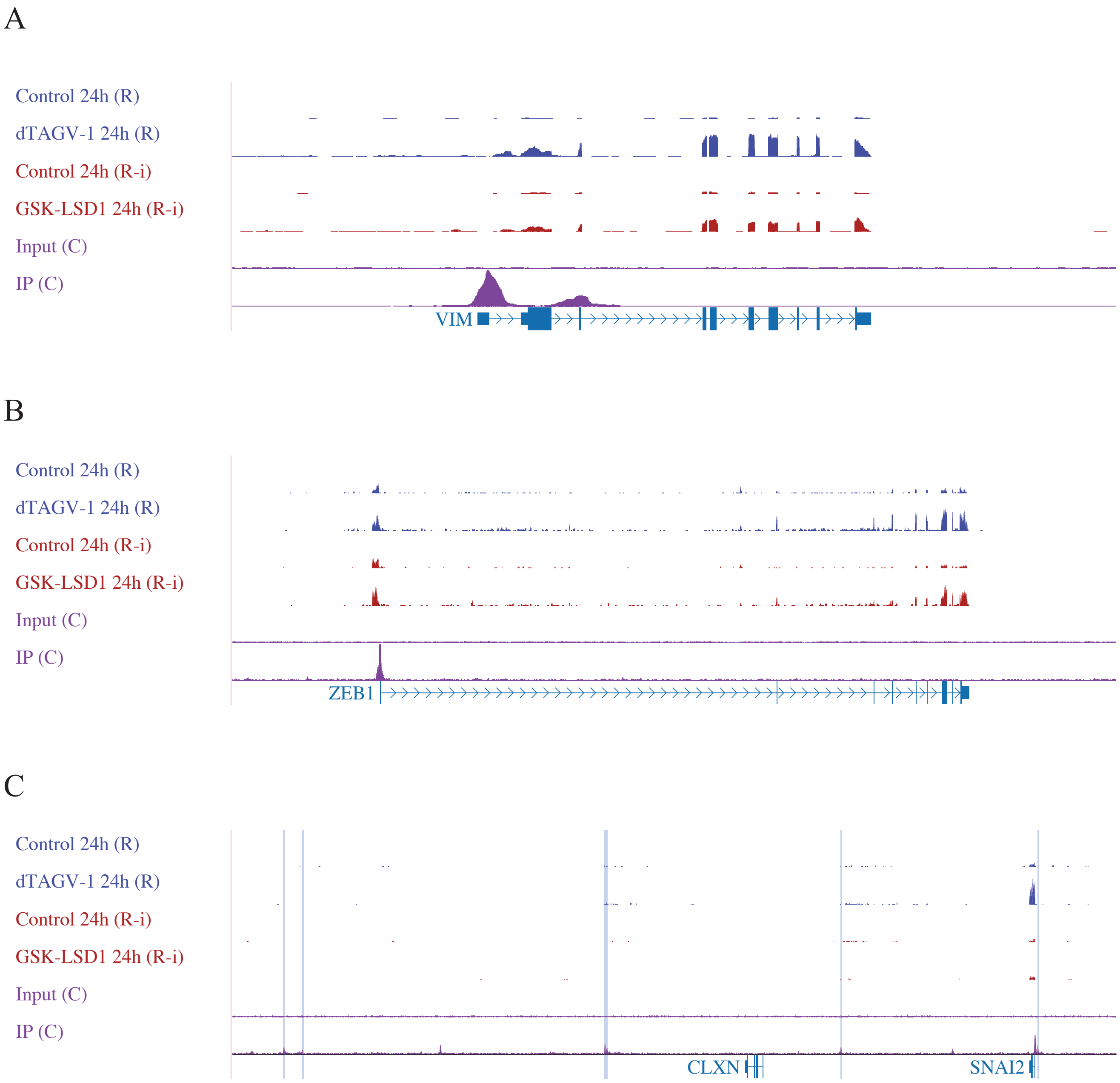


**Supplementary Fig. 7** LSD1 regulated genes differently when bound to enhancers vs promoters. (A-C) Genome tracks showing LSD1 mRNA reads ollowing LSD1 depletion or inhibition and for the LSD1 binding at the core mesenchymal factors *VIM* (A), *ZEB1* (B), and *SNAI2* (C). Known SNAI2 enhancer regions are highlighted in blue.
